# Sustained visual priming effects can emerge from attentional oscillation and temporal expectation

**DOI:** 10.1101/604702

**Authors:** Muzhi Wang, Yan Huang, Huan Luo, Hang Zhang

## Abstract

Priming refers to the influence that a previously encountered object exerts on future responses to similar objects. For many years, visual priming has been known as a facilitation and sometimes an inhibition effect that lasts for an extended period of time. It contrasts with the recent finding of an oscillated priming effect where facilitation and inhibition alternate over time periodically. Here we developed a computational model of visual priming that combines rhythmic sampling of the environment (attentional oscillation) with active preparation for future events (temporal expectation). Counterintuitively, it shows both the sustained and oscillated priming effects can emerge from an interaction between attentional oscillation and temporal expectation. The interaction also leads to novel predictions such as the change of visual priming effects with temporal expectation and attentional oscillation. Reanalysis of two published datasets and the results of two new experiments of visual priming tasks with male and female human participants provide support for the model’s relevance to human behavior. More generally, our model offers a new perspective that may unify the increasing findings of behavioral and neural oscillations with the classic findings in visual perception and attention.

**Significance Statement:** There is increasing behavioral and neural evidence that visual attention is a periodic process that sequentially samples different alternatives in the theta frequency range. It contrasts with the classic findings of sustained facilitatory or inhibitory attention effects. How can an oscillatory perceptual process give rise to sustained attention effects? Here we make this connection by proposing a computational model for a “fruit fly” visual priming task and showing both the sustained and oscillated priming effects can have the same origin: an interaction between rhythmic sampling of the environment and active preparation for future events. One unique contribution of our model is to predict how temporal contexts affects priming. It also opens up the possibility of reinterpreting other attention-related classic phenomena.

## Introduction

Prior encounter with an object may speed up one’s response to a similar object, a pervasive phenomenon in human cognition known as priming (Schacter & Buckner, 1998; Tulving & Schacter, 1990). The visual priming task has proved to be a powerful behavioral paradigm in probing the temporal dynamics of human attention (Eimer & Schlaghecken, 2003). In its simplest version (Figure 1A), participants judge the pointing direction of a target arrow preceded by a congruent or incongruent prime. The priming effects—response time (RT) for incongruent trials minus RT for congruent trials—may vary with the prime-to-target delay (“SOA”). Classic findings include sustained positive priming in unmasked priming tasks (Neumann & Klotz, 1994) and transient positive priming followed by sustained negative priming in masked priming tasks (Eimer, 1999; Eimer & Schlaghecken, 1998).

**Figure 1.**
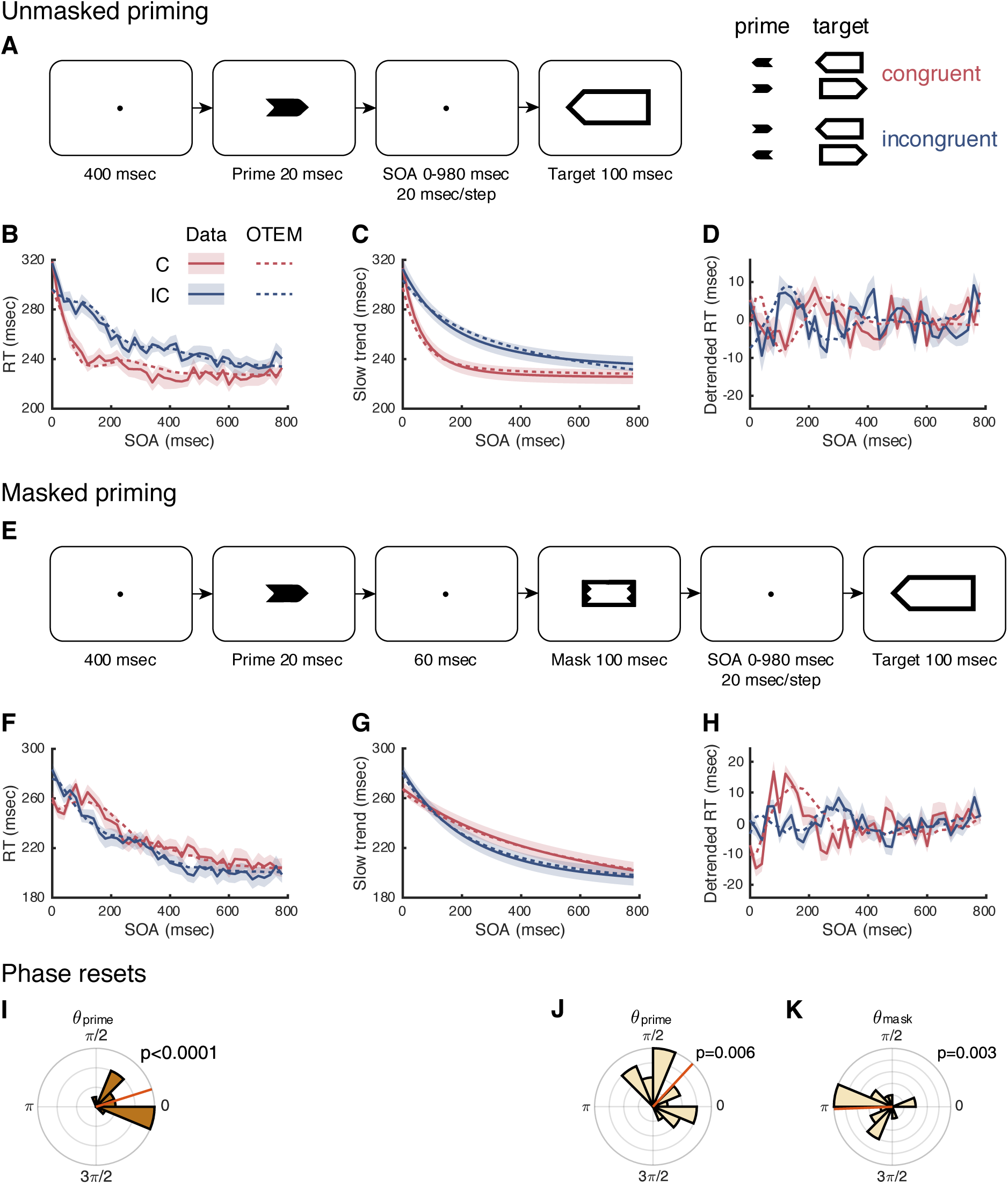
Oscillated Temporal Expectation Model (OTEM) reproduces the three priming effects in Huang et al. (2015). **A**, the unmasked priming task. Participants reported the pointing direction of the target arrow by key press. When the prime and the target pointed to the same direction, they were congruent (*C*), otherwise incongruent (*IC*). SOA was defined for the unmasked priming task as the delay between the onsets of the prime and the target. **B**, mean RT as a function of SOA in the unmasked priming task, separately for the congruent (red) and incongruent (blue). Solid lines denote data. Dotted lines denote OTEM fits. Shadings denote SEM. Following Huang et al. (2015), raw RTs of the unmasked priming task (**B**) were decomposed into slow trends (**C**) and detrended RTs (**D**), where positive priming (**C**) and oscillated priming (**D**) were observed. **E**, the masked priming task. The same as unmasked priming task except that a mask followed the prime and the delay between the onsets of the mask and the target was referred as SOA. **F**, mean RT as a function of SOA in the masked priming task, where positive priming followed by negative priming (slow trends, **G**) and oscillated priming (detrended RTs, **H**) were observed. **I**, Phase of attentional oscillation reset by the prime in the unmasked priming task. **J, K**, phases of attentional oscillation reset by the prime and the mask in the masked priming task. *θ*_prime_ and *θ*_mask_ were parameters of OTEM estimated from data. Rose plot shows the distribution of *θ*_prime_ or *θ*_mask_ across participants, which had significantly coherent values (Rayleigh tests, with *p* values marked on panels). The mean phase (red line) was close to 0 for *θ*_prime_ and to π for *θ*_mask_. That is, attention is biased towards the congruent in the next half cycle following the prime and towards the incongruent in the next half cycle following the mask.

Recently, a 3–5 Hz (close to theta-band) RT oscillation has been separated from these classic priming effects by spectral methods (Huang, Chen, & Luo, 2015): RT fluctuates up and down as a function of SOA in cycles of approximately 300 msec, with the peaks of congruent trials accompanied by the valleys of incongruent trials and vice versa (Figure 1DH). Such ongoing oscillations are visually pronounced even in the raw RTs of extensively tested individual subjects (Figure 4 of Huang et al., 2015). The finding of oscillated priming was a surprise but not entirely unexpected. On one hand, “sawtooth” patterned priming effects (Lingnau & Vorberg, 2005; Sumner & Brandwood, 2008) and oscillatory spatial cueing effects (Chen, Wang, Wang, Tang, & Zhang, 2017; Song, Meng, Chen, Zhou, & Luo, 2014) were reported. On the other hand, theta-band behavioral oscillations are widely observed in human perception (Benedetto and Morrone, 2017; Landau et al., 2015; Tomassini et al., 2015; Wutz et al., 2016; see VanRullen, 2016 for a review), which may reflect a rhythmic perceptual sampling of the environment.

**Figure 2.**
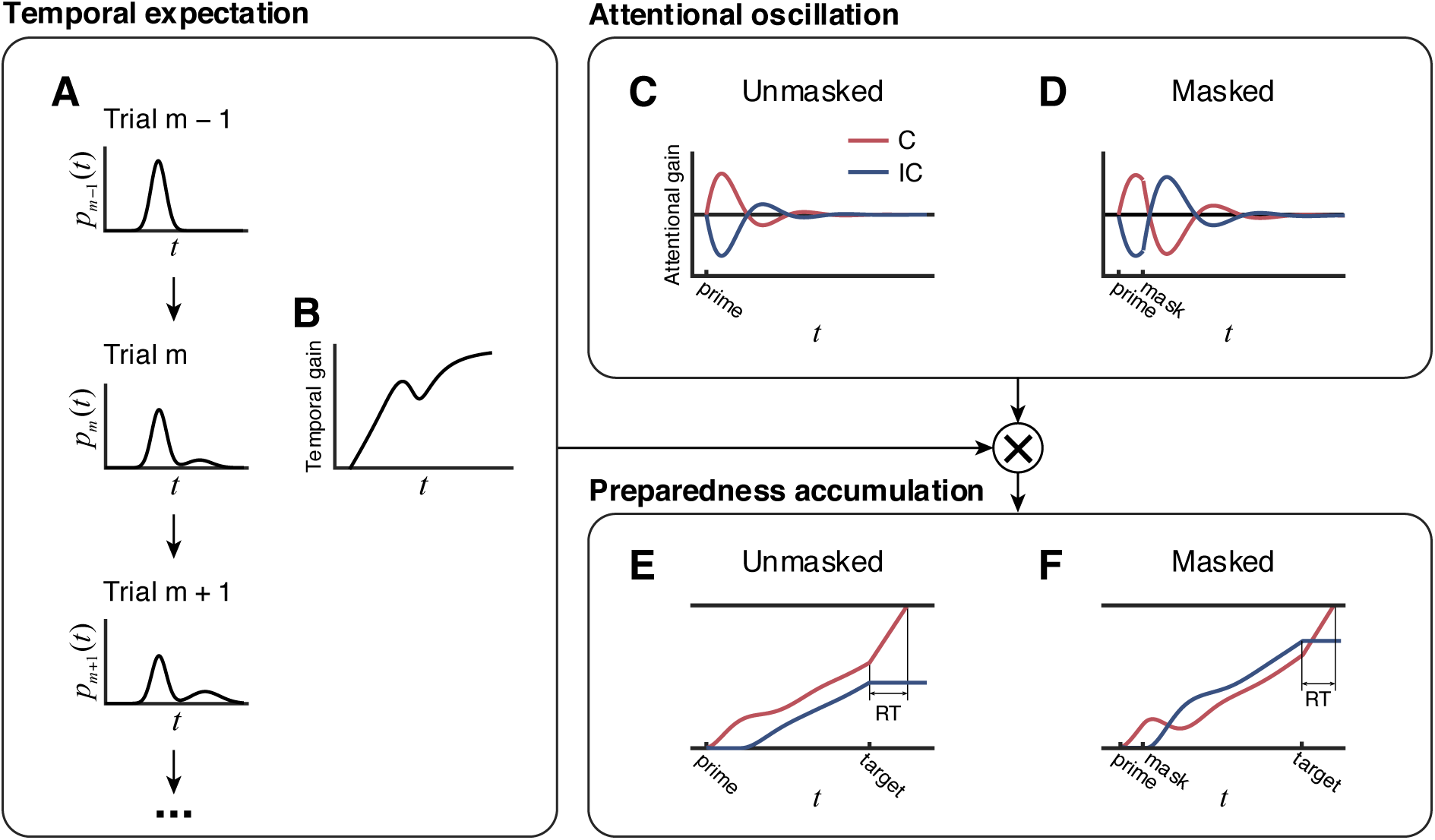
Illustration of the assumptions of OTEM. Temporal expectation: After each trial, the belief of SOA distribution (*p*_*m*_(*t*)) is updated following the delta rule (**A**). Within a specific trial, the temporal gain (**B**) is computed based on *p*_*m*_(*t*) and updated from moment to moment until target onset. **Attentional oscillation**: The attentional gain for the congruent (*C*, in red) and incongruent (*IC*, in blue) oscillates over time, whose amplitude and phase are reset by the prime and the mask (**C, D**). In the masked priming task, the oscillation induced by the mask superimposes on the oscillation induced by the prime. **Preparedness accumulation**: Before target onset, two threads of preparedness accumulate over time separately for the congruent and incongruent, with the accumulating speed at each moment determined by the product of the corresponding attentional and temporal gains (**E, F**). After target onset, only the thread for the realized target continues to accumulate until it reaches the responding bound. The RT for a specific target is thus determined by the preparedness accumulated for the target up to its onset. Intuitively, the attentional oscillation in the unmasked priming task (**C**) gives the congruent thread a head start in the race (**E**), thus leading to sustained positive priming. In contrast, the mask in the masked priming task resets attentional oscillation (**D**) and reverses the ranking of the congruent and incongruent threads (**F**), thus leading to transient positive priming followed by sustained negative priming.

**Figure 3.**
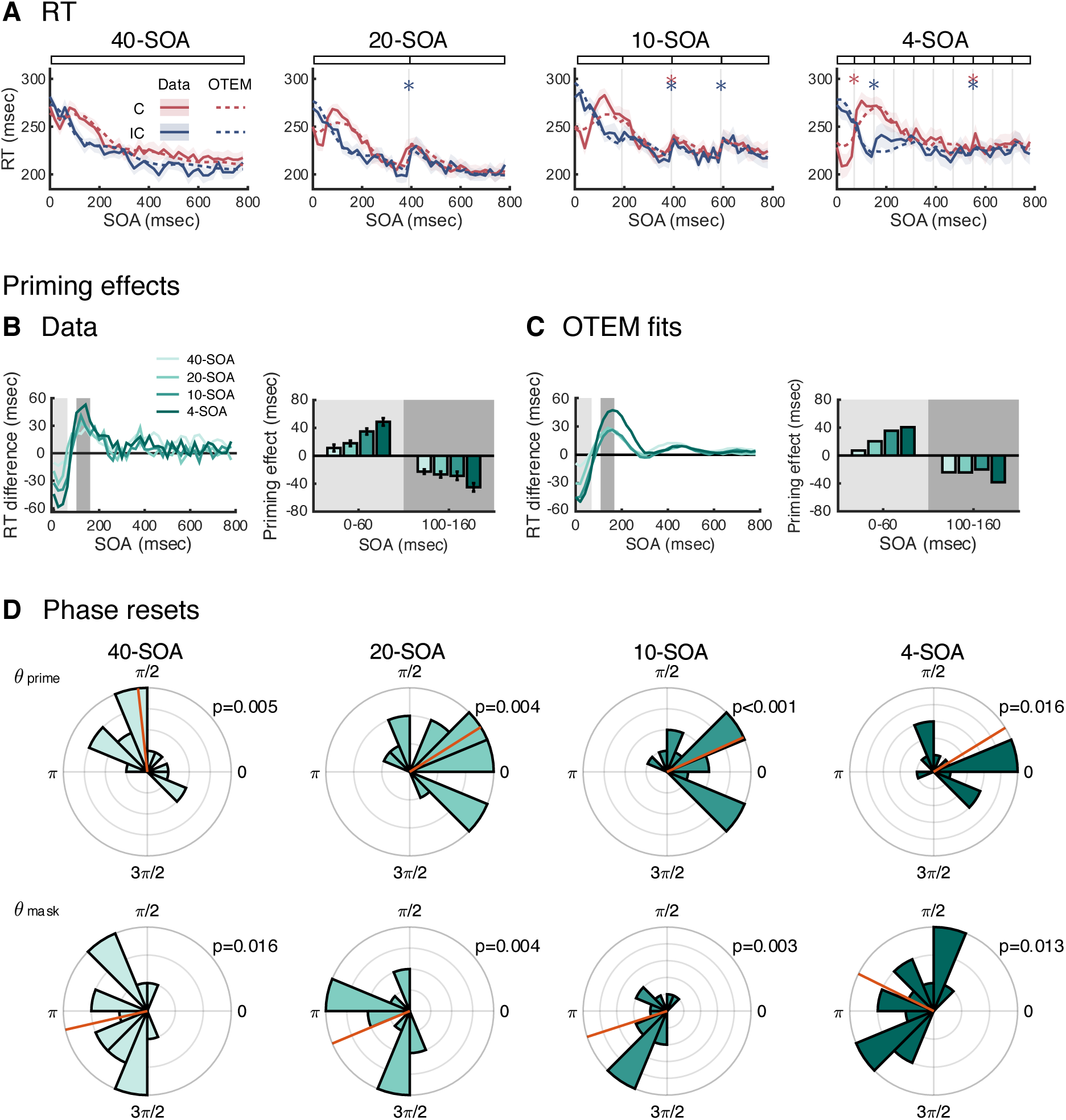
OTEM predicts the effects of temporal expectation in the temporal context experiment. **A**, mean RT as a function of SOA, separately for the congruent (red) and incongruent (blue). Solid lines denote data. Dotted lines denote OTEM fits. Shadings denote SEM. Each panel is for one temporal context condition, with the experimental manipulation schematized above the panel (see text). Asterisks indicate statistically significant “jumps” at block borders (*p* < 0.05). **B**, observed priming effects and **C**, OTEM predicted priming effects as functions of SOA. “RT difference” refers to the difference in RT between the congruent and incongruent. “Priming effect” was computed as the mean of incongruent RT minus congruent RT within a specific time window that is indicated by shading. Error bars denote SEM. Around the peaks of positive priming (SOA 0–60 msec, in light gray shading) and negative priming (SOA 100–160 msec, in dark gray shading), the magnitudes of mean priming effects (bar graphs) increased over the four conditions. **D**, phases of attentional oscillation reset by the prime and the mask. Conventions follow **Figure 1JK**. These phases resembled their counterparts of Huang et al. (2015) (see Figure **1JK**).

**Figure 4.**
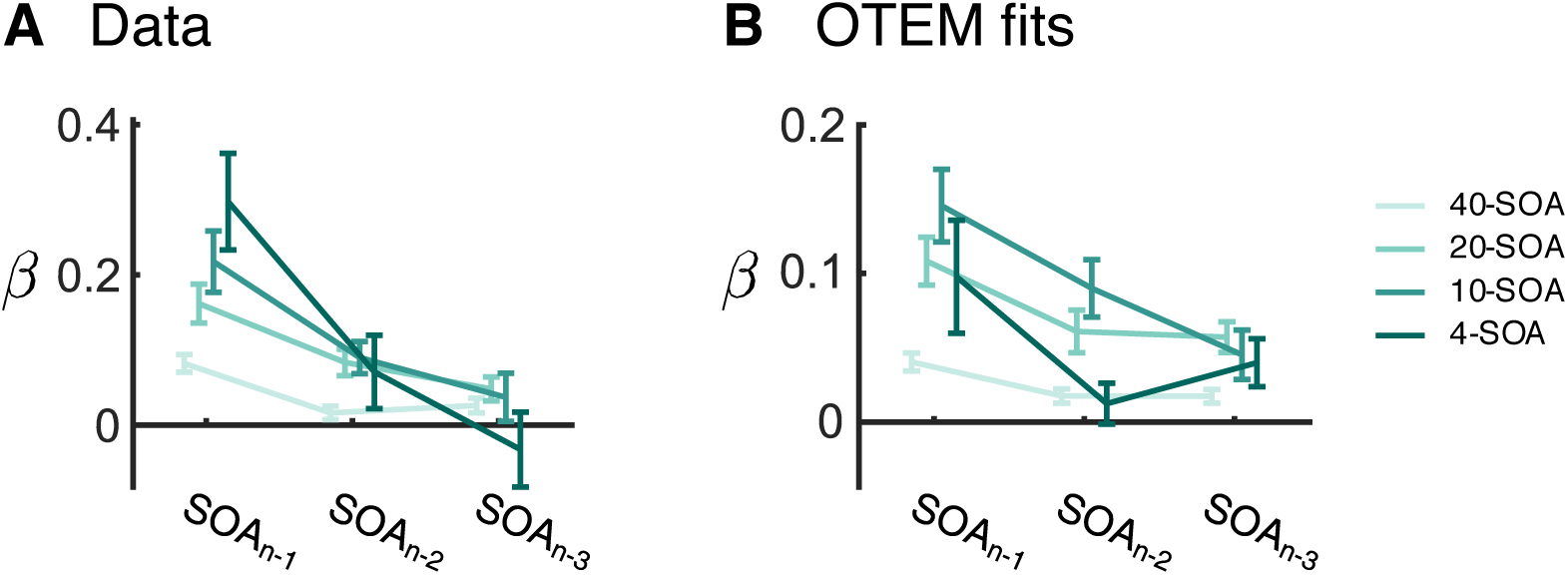
Sequential effects of SOAs. OTEM predicts that when the preceding trials have longer SOAs, the temporal expectation for the current SOA would be longer, thus the preparing speed would be lower, resulting in a longer RT. To quantify this possible sequential effect, we regressed the current RT over the SOAs of the last three trials, with the RTs of the last three trials and the current SOA serving as nuisance covariates. The regression coefficients (*β*) from OTEM fits (**B**) closely resembled the patterns of the data (**A**).

What may explain the coexistence of the classic and oscillated cueing/priming effects, which seem to suggest two conflicting time schedules for attention orienting? Previous theories on spatial cueing (Klein, 2000; Nobre & Rohenkohl, 2014) or visual priming (Bowman, Schlaghecken, & Eimer, 2006; Eimer & Schlaghecken, 2003; Huber, 2014; Lleras & Enns, 2004; Sumner, 2007) have been devoted to the classic effects, which apparently cannot predict any ongoing oscillations. To explain the oscillatory RT effects, an attentional oscillation mechanism seems to be necessary. An additional factor that may have a major impact on RT but is largely neglected by previous cueing/priming theories is temporal expectation. Even though the cue/prime is non-informative about the identity of the target, it signals the countdown to the target arrival. Temporal expectation has thus been assumed in the foreperiod task (Los, Kruijne, & Meeter, 2014; Los & Van Den Heuvel, 2001; Luce, 1986; Niemi & Näätänen, 1981; Nobre, Correa, & Coull, 2007) to account for the decrease of RT with increasing SOA and is likely to explain a similar RT effect in cueing/priming (e.g. Figure 1BF).

Here we proposed a computational model that introduces attentional oscillation and temporal expectation into the visual priming task to predict the RT patterns. Its basic idea is close to “active sensing” (Lakatos, Karmos, Mehta, Ulbert, & Schroeder, 2008; Lakatos et al., 2013) that the perceptual system actively predicts and prepares for potential future events (Nobre & van Ede, 2017; Summerfield & de Lange, 2014). Meanwhile, theta-band attentional oscillations provide fine-grained temporal windows of excitation and inhibition for the accumulation of preparedness (van Vugt, Simen, Nystrom, Holmes, & Cohen, 2012). We call the model Oscillated Temporal Expectation Model (OTEM), whose key assumptions are illustrated in Figure 2 and unfolded in the Results. In brief, OTEM predicts not only the oscillated priming effect and the decrease of RT with increasing SOA, but also the classic priming effects. That is, sustained positive and negative priming effects can arise as emergent properties of the model, despite that no sustained excitation or inhibition mechanisms are assumed.

## Materials and Methods

### Experimental Design and Statistical Analysis

We had analyzed data from two published experiments of Huang et al. (2015) and two new experiments (the temporal context experiment and the Int0-Int60 experiment). The experiments had been approved by the ethics committees of Institute of Biophysics at Chinese Academy of Sciences (#2006-IRB-005) and School of Psychological and Cognitive Sciences at Peking University (#2015-03-13c). All participants provided informed consent before the experiment. Participants were paid for their time and might also receive bonus for an accuracy of above 95%. The study was not preregistered. No statistical methods were used to predetermine sample sizes but our choice of sample size was based on previous work, including Huang et al. (2015).

#### Huang et al. (2015)

The unmasked and masked priming experiments of Huang et al. (2015) that we reanalyzed had 16 participants (6 male) and 18 participants (8 male) respectively. Participants were seated in a dark room, 57 cm from the computer monitor (refresh rate 100 Hz), with their heads stabilized by a chinrest. Stimuli were black visual shapes (0 cd/m^2^) presented on a gray background (0.99 cd/m^2^). As shown in Figure 1, the prime was a left- or right-pointing solid arrow (2.86° × 1.21°), the mask was a rectangular outer shape (3.43° × 1.79°) with an inner cut that fit the prime, and the target was a larger left- or right-pointing hollow arrow (6.21° × 2.29°).

The temporal course of one trial is shown in Figure 1A for the unmasked priming task and in Figure 1E for the masked priming task. After a 400-msec fixation point, a 20-msec prime stimulus was presented. It was followed by a 100-msec target (unmasked priming), or a 100-msec mask and a 100-msec target (masked priming). Participants were asked to respond as fast as possible (time limit: 1500 msec) whether the target arrow pointed to the left or right. Their responses were recorded by a parallel-port response keypad.

The inter-stimulus interval between the prime and the mask in the masked priming task was 60 msec. The SOA (stimulus onset asynchrony) was defined for the unmasked priming task as the delay between the onsets of the prime and the target and for the masked priming task as the delay between the onsets of the mask and the target. Fifty different SOAs were sampled for each experiment, in a 20-msec step from 0 to 980 msec. When the prime and the target in a trial pointed to the same direction, the trial was referred to as congruent (*C*), otherwise incongruent (*IC*). Each combination of SOA and congruency was repeated for 16 times, with all trials randomly interleaved.

#### Temporal context experiment

The temporal context experiment was designed to test the temporal expectation component of OTEM. Sixty-four young adults (29 male, aged 19–27) participated, with 16 participants randomly assigned to each of the four conditions of the experiment. The experiment used the masked priming task of Huang et al. (2015) but consisted of four experimental conditions that differed in local temporal contexts. In the 40-SOA condition, 40 different SOAs (0 to 780 msec in a 20-msec step) were repeated in a random order, as in Huang et al. (2015). In the 20-SOA condition, the same 40 SOAs were divided into two blocks—the first 20 SOAs and last 20 SOAs, so that the SOA was expected to be within 0–380 msec for the former and within 400–780 msec for the latter. Similarly, the 10-SOA condition had 4 blocks of 10 SOAs and the 4-SOA condition had 10 blocks of 4 SOAs. In all the conditions, each combination of SOA and congruency was repeated for 12 times and the order of the blocks was randomized for each participant.

#### Int0-Int60 experiment

The Int0-Int60 experiment was designed to test the attentional oscillation component of OTEM. Nineteen participants (9 male, aged18–25) completed the experiment. One additional participant quitted after the first session and was excluded. In the masked priming experiment of Huang et al. (2015), the inter-stimulus interval between the prime and the mask was always 60 msec. In the Int0-Int60 experiment, we used the same masked priming task (response time limit: 1000 msec) but manipulated the prime-to-mask interval to be 0 or 60 msec, referred respectively as the Int0 and Int60 conditions. In each condition, the mask-to-target SOAs ranged from 0 to 620 msec in a 20-msec step, with each combination of SOA and congruency repeated for 20 times, randomly interleaved. Each participant completed the Int0 and Int60 conditions in two different sessions on two separate days. The order of the two conditions was counterbalanced across participants.

#### Statistical Analysis

Time-out trials, trials with premature responses, and trials with response times (RTs) outside the 3 SDs of each participant were excluded. For the unmasked and masked experiments of Huang et al. (2015), as in the original paper, SOAs longer than 800 msec were truncated before further statistical analysis. We followed the approach of Huang et al. (2015), decomposing raw RTs into slow trends and detrended RTs. For each participant and separately for the congruent and incongruent trials, we smoothed raw RTs using exponential fits to obtain slow trends. Subtracting slow trends from raw RTs resulted in detrended RTs.

Fast Fourier Transform (FFT, implemented by MATLAB function *fft*) was used for spectral analysis. FFT was performed on the detrended RTs of each participant and then averaged across participants in the frequency domain. The permutation procedure and multiple-comparison correction method of Huang et al. (2015) were performed to test the statistical significance of the amplitude spectrum of detrended RTs as follows, separately for the congruent and incongruent trials. We generated surrogate signals by randomly shuffling the detrended RTs across SOAs within each participant, applied FFT to these surrogate signals, and computed the mean amplitude spectrum across participants. This procedure was repeated for 1,000 times to produce a distribution of mean amplitude at each frequency point, from which we elicited the uncorrected *p* < 0.05 threshold for false positives. To correct for multiple comparisons, we set the maximum of the uncorrected thresholds across frequency points as the corrected *p* < 0.05 threshold (Nichols & Holmes, 2002; Song et al., 2014).

As in Huang et al. (2015), the 3–5 Hz phase difference between the congruent and incongruent was calculated for each participant as the mean phase difference across 3–5 Hz. Rayleigh tests were used to test the phase coherence across participants. Statistical tests and averaging on circular variables were implemented by Matlab toolbox CircStat (Berens, 2009).

### Oscillated temporal expectation model (OTEM)

See Results for a brief summary of the assumptions and motivations of OTEM.

#### Preparedness accumulation

We assume that two threads of preparedness are accumulated in parallel for the two potential targets after the prime and before target onset, each of which can be formulated as:

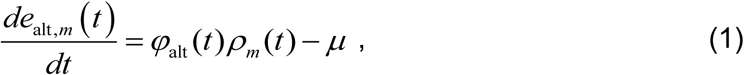

where `alt` denotes whether the potential target is congruent (C) or incongruent (IC) with the prime, *m* denotes trial number, *t* denotes the delay since prime, *e*_*alt,m*_(*t*) denotes accumulated preparedness, *φ*_*alt*_(*t*) denotes attentional gain, *ρ*_*m*_(*t*) denotes temporal gain, *µ* is a leaky rate parameter.

We assume *e*_alt,*m*_(0) = 0. To avoid premature responses before target onset, an upper boundary is imposed on *e*_alt,*m*_(0) = 0. After the target arrives, the preparedness for the target continues to accumulate at a constant rate *v* while the other thread stops. The expected RT for Trial *m* is the duration from target onset (*τ*_target,*m*_) to the moment the preparedness hits the response threshold *b* :

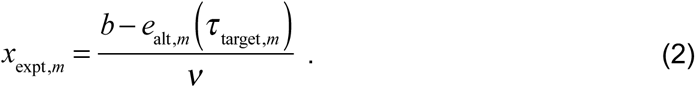

The observed RT *x*_obs,*m*_ is modeled by adding a log-normal noise:

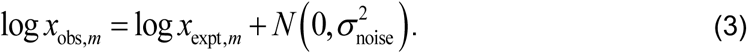

#### Attentional gain

We assume that the attentional gains for the two alternative targets add up to 2 and oscillate anti-phased over time, with the amplitude of oscillation decaying exponentially with time and the phase of oscillation reset by the onset of the prime or mask (Figure 2CD). We consider two alternative hypotheses about phase resets (Ding, Simon, Shamma, & David, 2016): substitutive and additive, that is, whether the attentional oscillation triggered by the mask overwrites or superimposes on that of the prime. Define

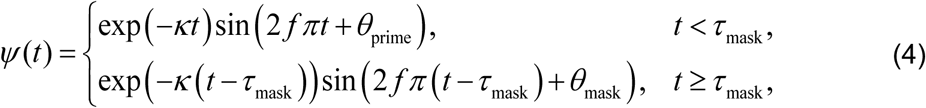

for substitutive phase reset and

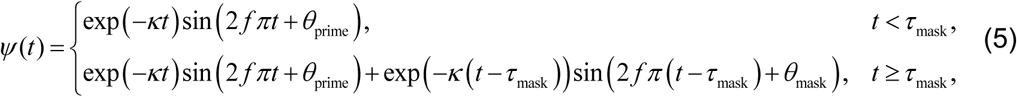

for additive phase reset, where *κ* is a free parameter for decaying rate, *f* denotes the frequency of attentional oscillation, *θ*_prime_ and *θ*_mask_ are free parameters for phases reset by the prime and mask. The moments *t* = 0 and *t* = *τ*_mask_ respectively correspond to the onsets of the prime and the mask. For the unmasked priming task, *τ*_mask_ → ∞. In our major model fitting procedures, we set *f* = 3.3 following the finding that oscillated priming peaks at 3.3 Hz (Huang et al., 2015).

The attentional gains for C and IC can be written as

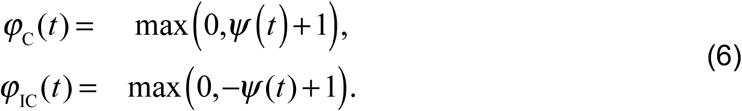

#### Temporal gain: trial-by-trial learning

We assume that the participant’s temporal expectation for the target is learned from previous experience trial by trial (Figure 2A) following the delta-rule learning (Rescorla & Wagner, 1972):

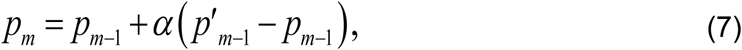

where *p*_*m*_ and *p*_*m*−1_ denote the expected distribution of *τ* _target_ at the beginning of Trial *m* and *m* − 1, 0 ≤ *α* ≤ 1 is a free parameter for learning rate. The 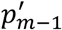 denotes the perceived distribution of *τ*_target_ on Trial *m* − 1, which is a Gaussian distribution around *τ* _target,*m*−1_:

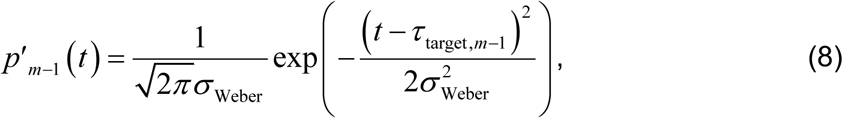

where *σ*_Weber_ follows Weber’s law, *σ*_Weber_ = *kτ*_target,*m*_ for unmasked and *σ*_Weber_ = *k*(*τ*_target,*m*_ − *τ*_mask_) for masked priming tasks, with Weber’s fraction *k* = 0.13 (Jazayeri & Shadlen, 2010). We set *p*_1_ to be a uniform distribution between the minimum and maximum *τ*_target_.

#### Temporal gain: real-time updating within trial

We assume that within each trial the brain keeps updating its expectation for the arrival time of the target (Figure 2B). On Trial *m*, the expected distribution of *τ*_target_ at Time *t* (*t* < *τ*_target,*m*_) is

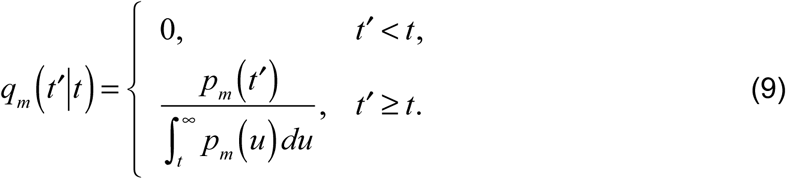

As a special case, the hazard rate of target onset at Time *t* is *h*(*t*) = *q*_*m*_ (*t*′ = *t*|*t*).

Treating the target as a future reward, we define the temporal gain at Time *t* as the expectation of the discounted future reward:

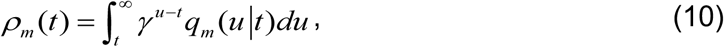

where 0 ≤ *γ* ≤ 1 is a free parameter for discounting rate.

In total, OTEM has nine parameters for unmasked priming tasks: learning rate, *α*; temporal discounting rate, *γ*; leaky rate, *µ*; decaying rate of oscillation, *κ*; phase reset by the prime, *θ*_prime_; maximum preparedness before target onset, *ζ*; accumulating rate after target onset, *v*; responding threshold, *b*; standard deviation of log-normal noise, *σ*_noise_. For masked priming tasks, there is one more parameter: phase reset by the mask, *θ*_mask_.

#### Extension of OTEM with a decision process

To explain the patterned errors in visual priming tasks, we enhanced OTEM with an error-prone decision process. The extended OTEM assumes that after target onset both the congruent and incongruent threads continue to accumulate, inhibiting each other and racing to the responding bound. The racing follows a noise-free version of the leaky competing accumulator model (Usher & McClelland, 2001):

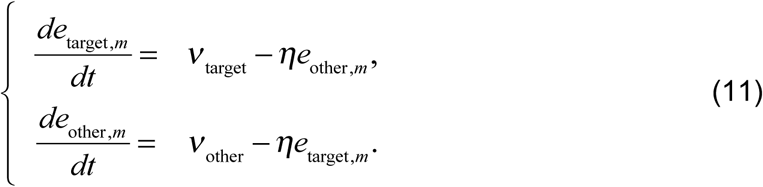

where *η* is a free parameter, *ν* _target_ and *ν*_other_ are accumulating rates for the target and the other threads. The values of *ν* _target_ and *ν*_other_ on each trial are randomly sampled from Gaussian distributions 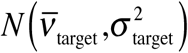 and 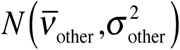, with 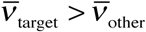.

Due to computational difficulties, we did not fit the extended OTEM and focused on qualitative predictions. To simulate for the results of Vorberg et al. (2003), we used the fitted parameters from the 4-SOA condition and manually set the additional parameters for the decision process.

### Oscillated urgency model

The oscillated urgency model is the same as OTEM, except that the preparedness for a potential target is not accumulated over time but determined by hazard rate. That is, Eq. 1 is replaced by

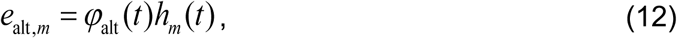

where *φ*_alt_(*t*) denotes attentional gain, *h*_*m*_(*t*) denotes hazard rate.

In total, oscillated urgency model has 7 free parameters: learning rate, *α*; decaying rate of oscillation, *κ*; phase reset by the prime or mask, *θ*; maximum preparedness before target onset, *ζ*; accumulating rate after target onset, *v*; response threshold, *b*; standard deviation of log-normal noise, *σ*_noise_.

### Constant accumulation model

The constant accumulation model is based on the accumulator model of Vorberg et al. (2003). The model assumes two accumulators for the two potential targets, the congruent and incongruent. Response is made when the absolute difference between the two accumulators exceeds the response threshold *b*. The same log-normal noise applies as in OTEM (Eq. 3).

Each accumulator follows

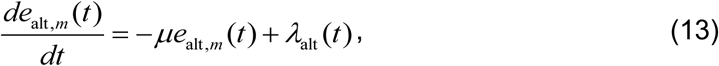

where *μ* is a free parameter for growth-decay rate, *λ*_alt_(*t*) denotes the rate of accumulation triggered by the prime, mask, or target as defined below. Following Vorberg et al. (2003), we assume that the prime and target would trigger positive accumulation rates for accumulators in their directions. To account for the observed effect of the mask (i.e. negative priming in masked priming tasks), we further assume that the mask would trigger a negative accumulation rate for the accumulator in the direction of the prime. That is, when the target is congruent with the prime,

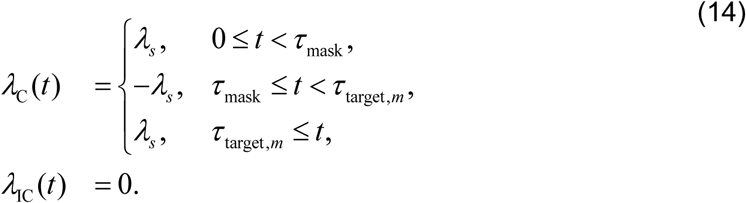

When the target is incongruent with the prime,

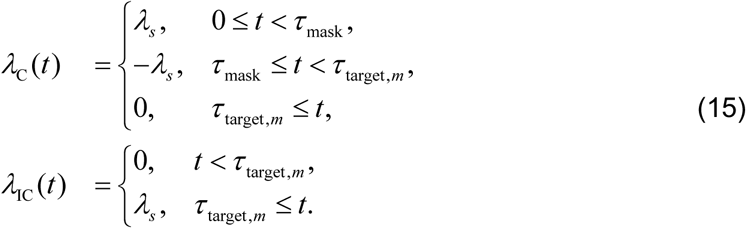

where *λ*_*s*_ is a free parameter, *t* = 0 corresponds to prime onset and *τ*_mask_ to mask onset. For the unmasked condition, *τ*_mask_ = *τ*_target,*m*_.

The threshold parameter *b* is redundant and fixed as 1. In total, the model has 4 free parameters: effect of stimulus, *λ*_*s*_; growth-decay rate, *µ*; maximum preparedness before target, *ζ*; standard deviation of log-normal noise, *σ*_noise_.

### Model fitting and comparison procedures

All the models were fitted on the individual level using the maximum likelihood estimate. For the Int0-Int60 experiment, where each participant completed two different prime-mask interval conditions, the same set of parameters were shared across the two conditions except for the responding threshold *b* and the standard deviation of log-normal noise *σ*_noise_. We used the interior-point algorithm of the *fmincon* function in MATLAB (MathWorks) to find the parameters that minimized negative log likelihood. To verify that we had found the global minimum, we repeated the search process with different starting points.

We used AICc—the Akaike information criterion with a correction for finite sample size (Akaike, 1974; Hurvich & Tsai, 1989)—for model comparison and calculated the protected exceedance probability of the group-level Bayesian model selection (Rigoux, Stephan, Friston, & Daunizeau, 2014; Stephan, Penny, Daunizeau, Moran, & Friston, 2009) as an omnibus measure across participants.

### Model simulation procedures

For each model and participant, 100 simulated datasets were generated based on the fitted parameters. These simulated datasets were analyzed in the same way as the real datasets, whose results were further averaged across participants to produce the model predictions on the group level.

### Data availability

All data and codes are available for download at https://osf.io/69by5/.

## Results

We tested OTEM quantitatively by first fitting it to the behavioral datasets of the unmasked and masked priming experiments in Huang et al. (2015) and found that it well reproduced all the priming effects as well as the overall RT patterns. Next, we performed two new experiments where temporal expectation or attentional oscillation was manipulated to test the key predictions that distinguish OTEM from previous accounts of visual priming effects (e.g. Bowman et al., 2006; Boy & Sumner, 2010; Vorberg et al., 2003). As predicted, the visual priming effects were influenced not only by SOA but also by temporal expectation and attentional oscillation. Third, we tested two alternative models and found neither of them could reproduce all the priming effects. Last, we showed that OTEM can be extended to predict error rates and atypical priming effects in the literature.

### Oscillated Temporal Expectation Model (OTEM)

The key assumptions of OTEM (Figure 2) is specified below (see Materials and Methods for details). For fluency of presentation, we only briefly describe their motivations here and leave a full treatment of the justification and implications of the assumptions to the Discussion.

#### Preparedness accumulation

We assume that responses are generated by an accumulation-to-bound process analogous to the racing models of perceptual decision-making (Luce, 1986). After the onset of the prime, two threads of preparation are separately accumulated for the two prospective targets—the congruent and incongruent—until the target arrives (Cisek, 2007; Cisek & Kalaska, 2010), after which only the preparedness for the realized target continues to accumulate. A response occurs when the responding bound is reached. The response time for a specific target is thus determined by the preparedness accumulated for the target up to its onset.

#### Temporal expectation

We assume that a belief of SOA distribution is learned from past trials following the delta rule (Rescorla & Wagner, 1972). On each trial, the brain uses this distribution as its (probabilistic) temporal expectation for the SOA of the incoming target and, as time elapses, keeps updating the expectation of remaining time until the target appears (de Lange, Rahnev, Donner, & Lau, 2013; McGuire & Kable, 2013).

At any specific moment, the temporal expectation—a probability distribution over all future time points—decides how urgent the brain should prepare for prospective targets. Similar to previous studies on temporal expectation (Bueti, Bahrami, Walsh, & Rees, 2010; Janssen & Shadlen, 2005; Sharma et al., 2014), we also considered the participant’s temporal uncertainty and used Weber’s law (Allan, 2001; Gibbon, 1977) to model it, with the Weber fraction parameter set to be a constant estimated from previous research (Jazayeri & Shadlen, 2010).

To map the influence of the distribution of temporal expectations to a single value, we introduce an economic concept: temporal discounting—the longer the expected delay of a reward, the lower the reward is valued in the brain (Frederick, Loewenstein, & O’Donoghue, 2002; Kable & Glimcher, 2007). In our case, it implies that the expected probability of target onset at a time point in the farther future has a smaller influence on the current preparing rate. Analogous to the application of temporal discounting in the reinforcement learning literature (Sutton & Barto, 1998), we assume that the preparing rate at a specific moment is proportional to the *temporal gain* at the moment, defined as the expected sum of the temporally discounted values of all future probabilities.

#### Attentional oscillation

There is increasing evidence that visual attention is a periodic process that sequentially samples different alternatives in the theta frequency range (Fiebelkorn, Saalmann, & Kastner, 2013; Kienitz et al., 2018; Landau & Fries, 2012; Landau et al., 2015; Song et al., 2014; Tomassini et al., 2015; VanRullen, 2016). Spontaneous theta-band neuronal oscillations are fairly common in the human brain and are readily reset by task-relevant stimuli (Rizzuto et al., 2003; Tesche and Karhu, 2000; see Buzsaki, 2006 for a review). Further, the oscillatory entrainment of sensory stimuli is anti-phased for the attended and unattended stimuli (Lakatos et al., 2013).

Following these empirical findings, we assume that the focus of attention switches gradually between the two prospective targets in theta-band oscillations, leading to anti-phased attentional gains for the congruent and incongruent. The phase of the attentional oscillation is reset by the prime and the mask, while its amplitude decreases exponentially with time, as suggested by stimulus-induced resets of theta-band neuronal oscillations (Buzsáki, 2006; Makeig et al., 2002; McCartney, Johnson, Weil, & Givens, 2004; Rizzuto et al., 2003; Tesche & Karhu, 2000) and behavioral oscillations (Fiebelkorn et al., 2013; Landau & Fries, 2012; Song et al., 2014). In our modeling, we consider two alternative hypotheses about phase resets (Ding et al., 2016): substitutive and additive, that is, whether the attentional oscillation induced by the mask overwrites or superimposes on that of the prime. We assume that the preparing rate for the congruent or incongruent target is scaled by its attentional gain as well as by the temporal gain.

Buzsaki (2006) suggested that theta-band neuronal oscillations serve as synchronizers to integrate information over time. Analogously, the attentional oscillation assumed in OTEM can in effect help to integrate temporal expectations over time.

To avoid introducing too many degrees of freedom into OTEM, some of our assumptions were simplified. For example, we assumed a transition from oscillatory dynamics to constant preparedness accumulation for the realized target immediately after target onset. To assume a delayed transition would be more realistic, since it takes time to detect the target. Given that the target appears at central vision with high contrast, such a delay tends to be negligible and whether to include it or not should have little influence on our conclusions.

### OTEM reproduces both the classic and oscillated priming effects

The datasets of Huang et al. (2015) include one unmasked experiment (*N* = 16) and one masked priming experiment (*N* = 18), where RTs for the congruent and incongruent varied as functions of SOA (Figure 1). Fifty different SOAs were sampled for each experiment, in a 20-msec step from 0 to 980 msec. Decomposing each raw RT curve into a slow trend and a detrended curve, Huang et al. (2015) identified the classic and oscillated priming effects respectively in the two components.

We fit OTEM to the RTs of individual trials for correct responses for each participant using maximum likelihood estimates and plot the model fits against data (Figure 1BF, dotted vs. solid curves). We considered two versions of OTEM, differing in whether the phase reset of attentional oscillation is additive or substitutive. For the experiments of Huang et al. (2015) and for the temporal context experiment introduced below, the two OTEMs had similar fits to the data. For these experiments, we thus only present the results of the “additive” OTEM, which had a slightly better goodness-of-fit than the “substitutive” OTEM (Figure 6O). Unless explicitly specified, OTEM hereafter refers to the additive OTEM.

**Figure 5.**
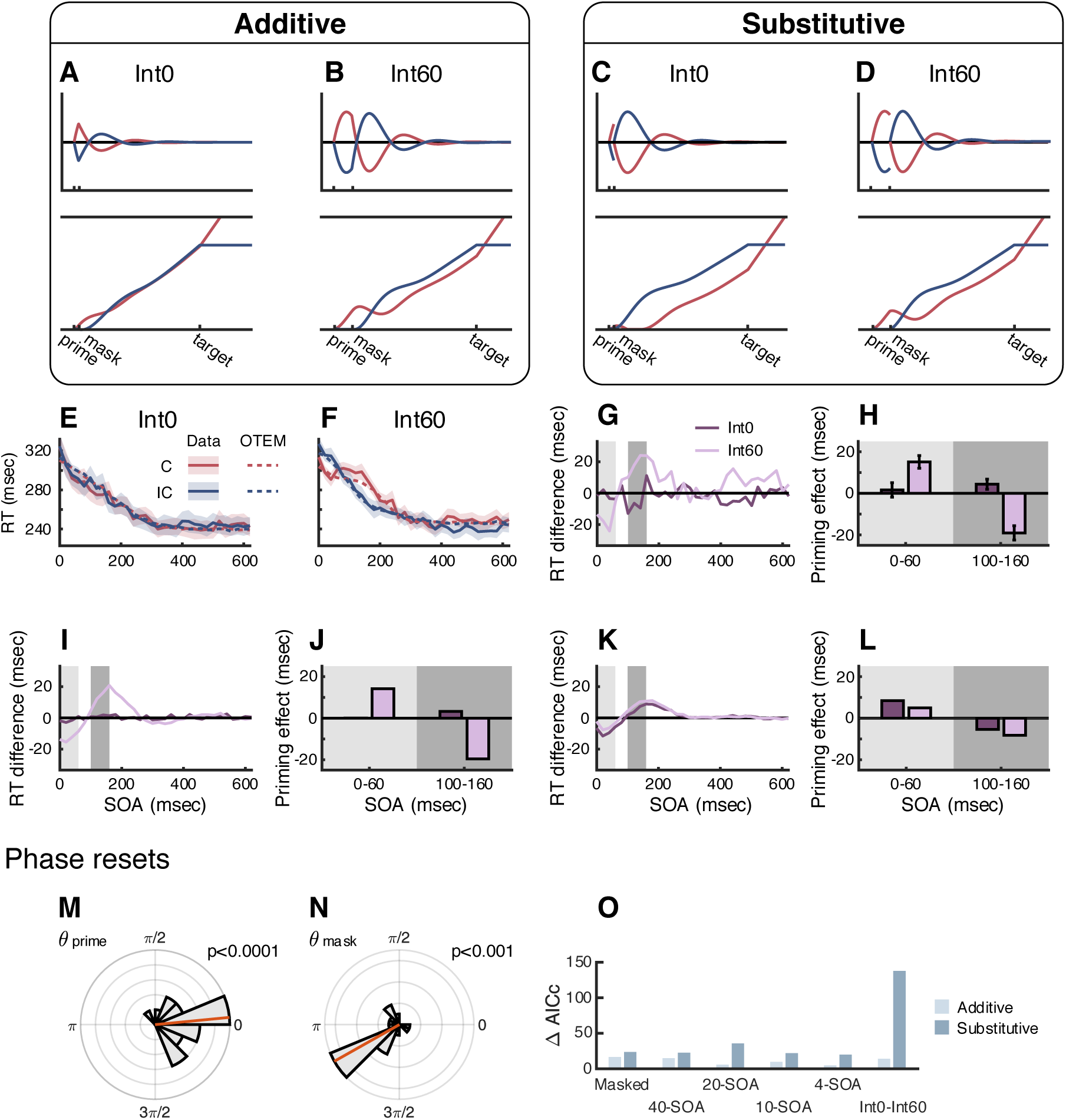
Additive OTEM predicts the vanish of priming effects at minimum prime-mask interval in the Int0-Int60 experiment. **A, B**, illustration of the additive OTEM predictions. **C, D**, illustration of the substitutive OTEM predictions. Conventions follow **Figure 2**. The prime and the mask are assumed to induce cycles of attentional oscillations that are anti-phased (i.e., *θ*_prime_ = 0 and *θ*_mask_ = π), respectively favoring the congruent and incongruent in the next half cycle. The assumptions of the additive and substitutive OTEM models differ in whether the attentional oscillation induced by the mask superimposes on or overwrites that of the prime. The two OTEM models have similar predictions for the Int60 condition but distinctively different predictions for the Int0 condition. **E, F**, mean RT as a function of SOA in the Int0 (**E**) and Int60 (**F**) conditions, separately for the congruent (red) and incongruent (blue). Solid lines denote data. Dotted lines denote additive OTEM fits. Shadings denote SEM. **G**, observed RT difference between the congruent and incongruent as a function of SOA for the Int0 (dark purple) and Int60 (light purple) conditions. Light gray shading and dark gray shading respectively denote the time windows of positive priming (0–60 msec) and negative priming (100–160 msec) defined earlier in the temporal context experiment. **H**, observed priming effects (incongruent RT minus congruent RT) averaged in the two time windows. In the Int60 condition, positive priming was followed by negative priming. In the Int0 condition, there were little priming effects in either time window. **I, J**, additive OTEM fits for priming effects, which replicated the observed lack of priming effects in the Int0 condition as well as the observed positive and negative priming effects in the Int60 condition. **K, L**, substitutive OTEM fits for priming effects, which failed to replicate the observed differences between the Int0 and Int60 conditions. **M, N**, phases of attentional oscillation reset by the prime and the mask. *θ*_prime_ and *θ*_mask_ were estimated phase parameters in the additive OTEM that were shared across the Int0 and Int60 conditions. Conventions follow **Figure 1I-K. O**, model comparisons between the additive OTEM and substitutive OTEM models for each experiment or condition. The unmasked masking experiment was not included because the two OTEM models are mathematically equivalent for unmasked priming. The summed ΔAICc across participants were plotted, with smaller ΔAICc indicating better goodness-of-fit. The additive OTEM outperformed the substitutive OTEM in all the experiments.

**Figure 6.**
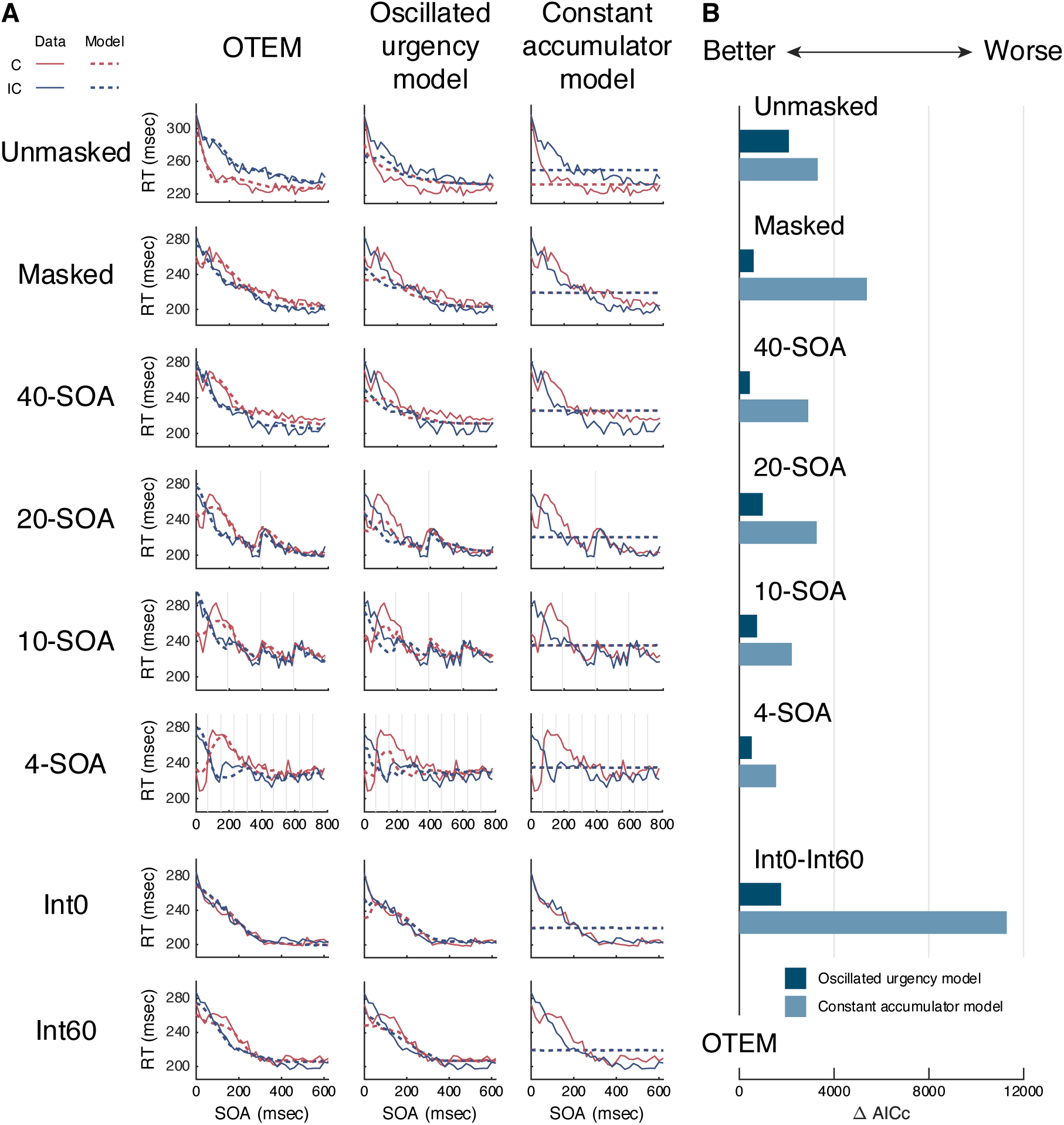
OTEM outperforms alternative models. **A**, model fits of OTEM (left column), oscillated urgency model (central column) and constant accumulation model (right column). Mean RT is plotted as a function of SOA separately for the congruent (red) and incongruent (blue). Solid lines denote data. Dotted lines denote model fits. Each row is for one experimental condition. “Unmasked” and “Masked” respectively refer to the unmasked and masked priming experiments of Huang et al. (2015). “40-SOA”, “20-SOA”, “10-SOA”, and “4-SOA” refer to the four temporal context conditions in the temporal context experiment. “Int0” and “Int60” refer to the two conditions in the Int0-Int60 experiment. The model fits of the two alternative models had obvious deviations from the RT data. **B**, the summed ΔAICc of the oscillated urgency model and constant accumulation model relative to that of OTEM (i.e., OTEM as the 0 baseline). A greater-than-0 ΔAICc indicates worse fit than OTEM. In all experiments and conditions, both alternative models fit the RT data much worse than OTEM did.

The OTEM fits captured the following patterns of the observed RT curves. First, the slow trends for both the congruent and incongruent trials decreased monotonically with SOA at a decreasing speed. Second, in the slow trends of the unmasked priming task (Figure 1C), positive priming (i.e. incongruent RTs above congruent RTs) persisted throughout all SOAs, with the difference between the incongruent and congruent RTs first increasing then decreasing. Third, in the slow trends of the masked priming task (Figure 1G), positive priming occurred for short SOAs (< 100 msec) and changed into negative priming (i.e. congruent RTs above incongruent RTs) for longer SOAs. Fourth, the detrended RT curves (Figure 1DH) oscillated at frequencies of 3–5 Hz and were anti-phased for the congruent and incongruent (see Figure 8 for details).

**Figure 7.**
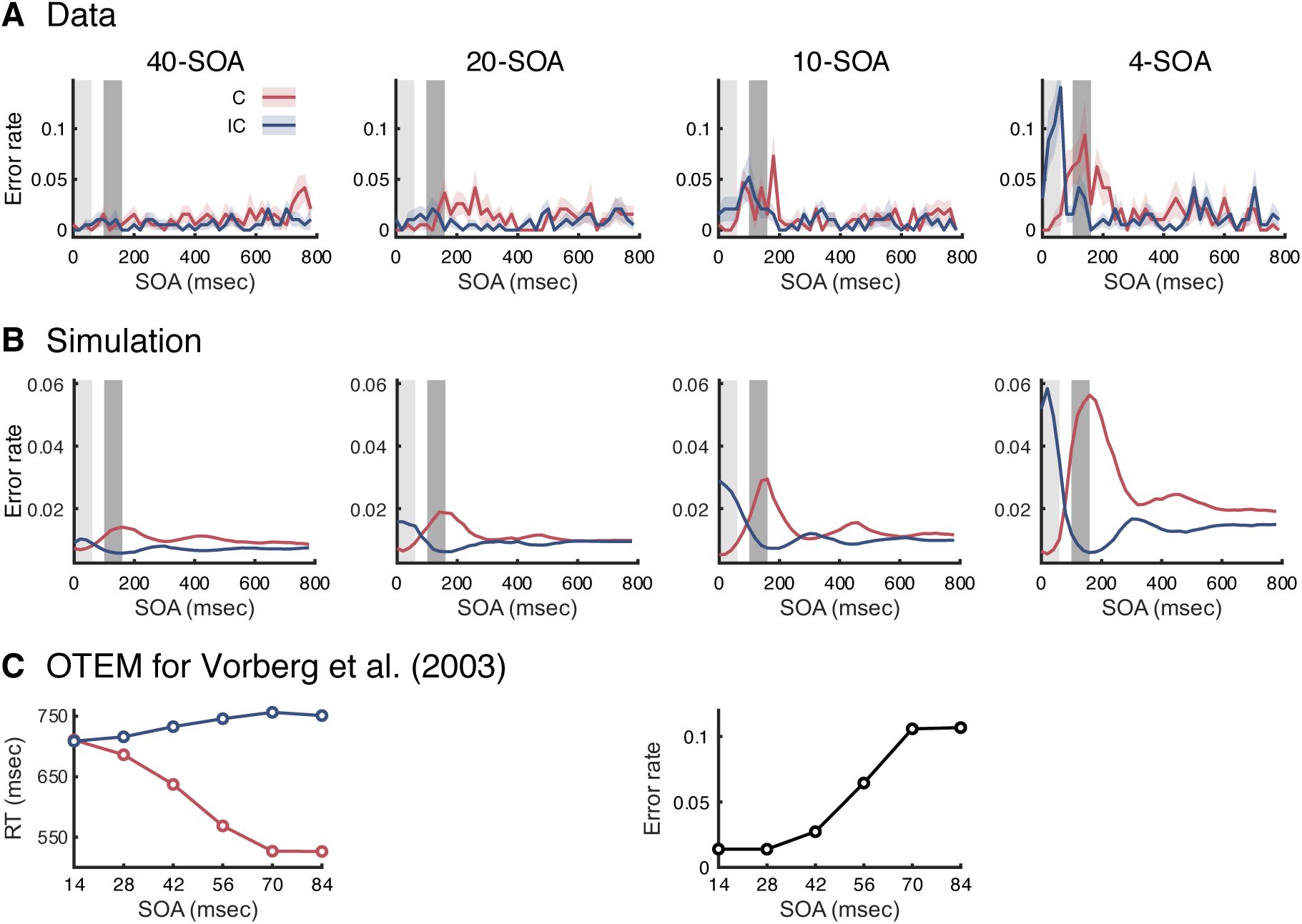
Extended OTEM explains error rates and atypical priming effects. **A**, mean error rate as a function of SOA, separately for the congruent (red) and incongruent (blue). Shadings around the mean error rates denote SEM. Light and dark gray shadings respectively denote the time intervals around the peak of positive and negative priming (defined as in **Figure 3**). Each panel is for one temporal context condition. **B**, mean error rate simulated by the extended OTEM, which reproduced the empirical pattern: positive priming was accompanied by a higher error rate for the incongruent and negative priming by a higher error rate for the congruent. **C**, mean RT and error rate simulated by the extended OTEM for the visual priming task of Vorberg et al. (2003), which captured the atypical patterns of their empirical findings: RTs for the congruent and incongruent respectively decreased and increased with SOA; error rates increased with SOA.

**Figure 8.**
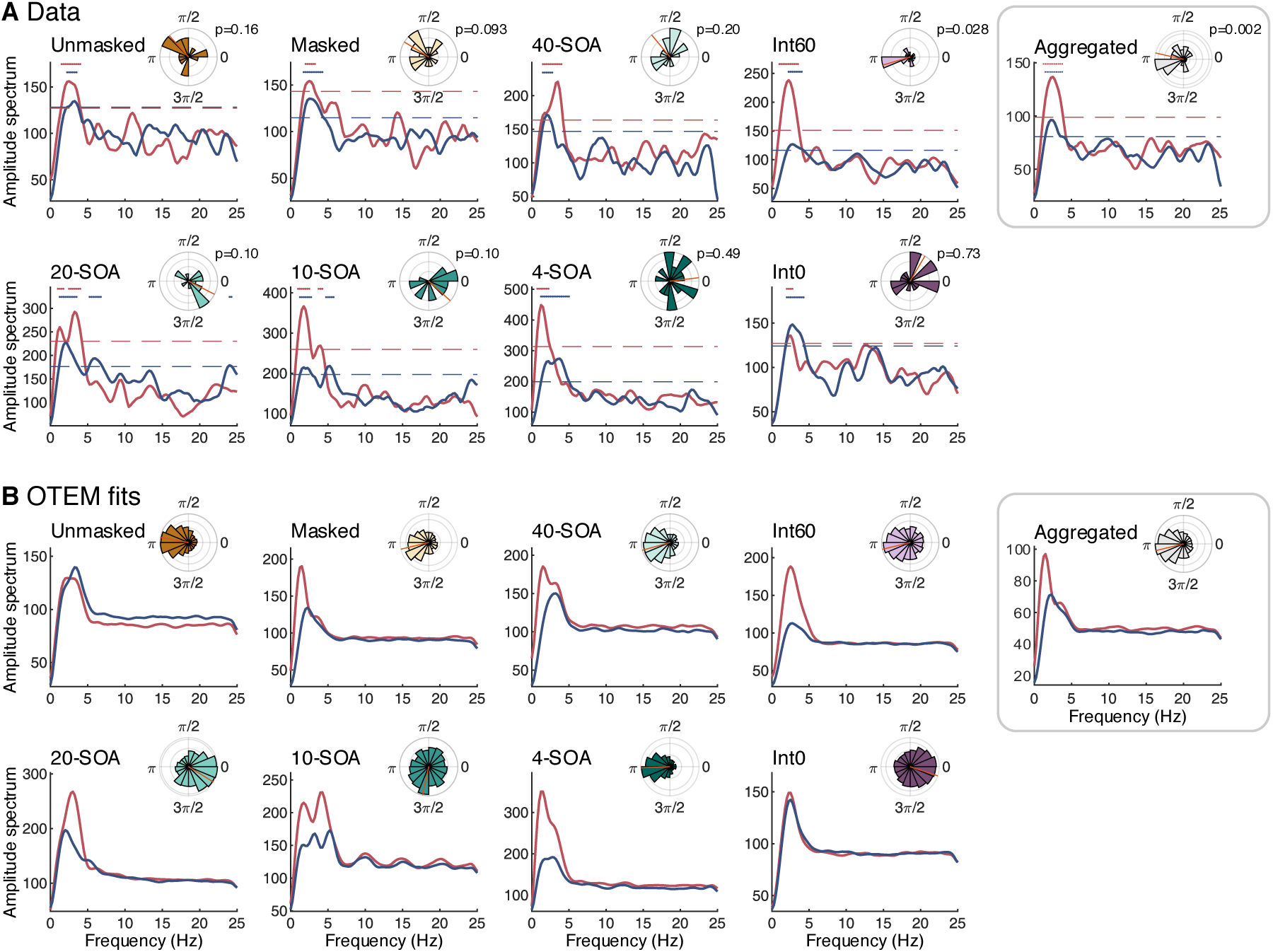
Spectral analysis of the detrended RT for data (A) and OTEM fits (B). Each panel denotes one experiment or condition. **Main plots**: amplitude spectrum, separately for the congruent (red curve) and incongruent (blue curve). Dash lines denote the 0.05 significance threshold of permutation test (corrected for multiple comparisons). Significant frequency points are marked by dots above the spectrum curves. **Insets**: phase difference between the congruent and incongruent in the 3–5 Hz oscillation. Rose plot shows the distribution of phase differences across participants, with the mean phase difference marked by a red line. The *p* value refers to the result of Rayleigh test on coherence. “Unmasked” and “Masked” respectively refer to the unmasked and masked priming experiments of Huang et al. (2015). “40-SOA”, “20-SOA”, “10-SOA”, and “4-SOA” refer to the four temporal context conditions in the temporal context experiment. “Int0” and “Int60” refer to the two conditions in the Int0-Int60 experiment. “Aggregated” refers to the results pooled across the similar conditions in three experiments: the masked, 40-SOA, and Int60 conditions. For all the experiments and conditions, the oscillations in the data were well captured by the OTEM fits.

In other words, OTEM reproduces both the classic and oscillated priming effects. Given that theta-band attentional oscillations are assumed in OTEM, it is probably not surprising the model can predict the observed oscillated priming. However, the classic positive and negative priming are emergent properties of the model, since no persistent attentional biases towards the congruent or incongruent have been assumed. That is, these classic priming effects arise also from the attentional oscillations. According to the phase parameters estimated from data (Figure 1I-K), both the prime and the mask induced phase resets that were coherent across participants (Rayleigh tests: unmasked prime, *r* = 0.82, *z* = 10.87, *p* < 0.0001; masked prime, *r* = 0.52, *z* = 4.84, *p* = 0.006; masked mask, r = 0.55, *z* = 5.47, *p* = 0.003). On average, the prime reset the phases of the attentional gains for the congruent and incongruent oppositely to approximately 0 and π (in sine), so that the preparedness of the congruent would accumulate faster than the incongruent in the first half cycle. Conversely, the mask reset the congruent and incongruent phases to approximately π and 0, giving the incongruent an advantage in the subsequent half cycle.

Here is an intuition of how the classic priming effects emerge from OTEM (see Figure 2 for illustrations): After the attentional oscillations are reset by the prime, the specific phase difference allows the attentional gain of the congruent to climb to the peak earlier than that of the incongruent, giving the congruent a lead in preparedness. For unmasked priming tasks, the congruent keeps the lead throughout the race despite that the congruent and incongruent receive alternating advantages in attentional gains, and thus positive priming is observed. For masked priming tasks, a second reset of the attentional oscillations by the mask reverses the lead, resulting in early positive priming and late negative priming.

According to our simulations, the emergence of the classic priming effects from OTEM in the experimental settings of Huang et al. (2015) is insensitive to the choice of most parameters in OTEM. Except for the phase reset parameters, changing the parameters of OTEM would only alter the overall RT and the magnitudes of the priming effects but hardly the qualitative finding of the priming effects.

### OTEM predicts the effects of temporal expectation

OTEM predicts that the RT of a trial should depend not only on its SOA (i.e. the actual moment of target onset), but also on the temporal expectation that is shaped by past trials. We tested this prediction in a new masked priming experiment, referred to as the temporal context experiment, which consisted of four experimental conditions that used the same set of SOAs but differed in local temporal contexts. In the 40-SOA condition, 40 different SOAs (0 to 780 msec in a 20-msec step) were repeated in a random order, as in Huang et al. (2015). In the 20-SOA condition, the same 40 SOAs were divided into two blocks—the first 20 SOAs and last 20 SOAs, so that the SOA was expected to be within 0–380 msec for the former and within 400–780 msec for the latter. Similarly, the 10-SOA condition had 4 blocks of 10 SOAs and the 4-SOA condition had 10 blocks of 4 SOAs. In all the conditions, each SOA was repeated for 24 times (12 congruent + 12 incongruent) and the order of the blocks was randomized for each participant. In the same procedure applied to the datasets of Huang et al. (2015), we fit OTEM to the RTs of individual trials for each participant in each condition and plot model fits against data (Figure 3A).

We found distinctive RT patterns in the four conditions (*N* = 16 for each condition). The 40-SOA condition replicated the results of the masked priming experiment of Huang et al. (2015), which would serve as a baseline. A hallmark of the other three conditions was a “jump” in RT at the border of two adjacent blocks, for example, the dramatic increase of RT from SOA=380 msec to SOA=400 msec in the 20-SOA condition. Out of the 26 block borders (separately for the congruent and incongruent) of the 20-SOA, 10-SOA, and 4-SOA conditions, 8 had significant RT increases (one-tailed *t*-tests, *p*s < 0.05). To assess the base rate of false positive increasing RTs, we performed *t*-tests for the same SOA pairs in the 40-SOA condition and had only one significant RT increase out of the 22 comparisons. The proportion of jumps at the block borders was significantly above the base rate (8/26 vs. 1/22, Fisher’s exact test, *p* = 0.028).

The OTEM fits agreed well with the observed RT patterns (Figure 3A). In particular, OTEM could predict the dramatic increase of RT at the borders of SOA blocks, although it makes no special assumptions about SOA blocks. According to OTEM, the closer the time approaches the last SOA of the block, the sooner the target is expected to arrive, the faster the accumulation of preparedness. Thus, the last SOA of a block could have a faster RT than the first SOA of its next block, despite the general decreasing trend of RT with SOA due to preparedness accumulation.

The magnitudes of priming effects differed across the four conditions (Figure 3B). Around the peak of positive priming (SOA = 0–60 msec), the mean priming effect (RT difference between the incongruent and congruent) went larger and larger from 40-SOA, 20-SOA, 10-SOA to 4-SOA (one-way ANOVA, *F*(3,60) = 13.02, *p* < 0.001). The mean peaked negative priming effect (SOA = 100–160 msec) exhibited a similar pattern (*F*(3,60) = 3.81, *p* = 0.014). Both patterns were captured by OTEM fits (Figure 3C).

The estimated phase reset parameters in OTEM (Figure 3D) for the 20-, 10-, and 4-SOA conditions were indistinguishable from those of the 40-SOA condition and the masked priming experiment of Huang et al. (2015) pooled (Watson-Williams test, *F*(3,78) = 2.53, *p* = 0.063 for prime phase; *F*(3,78) = 1.65, *p* = 0.18 for mask phase). That is, the attentional oscillations were little influenced by the temporal context of the experiment.

As we mentioned earlier, OTEM correctly predicts that the different temporal expectations associated with different contextual blocks may lead to RT “jumps” at the block borders (Figure 3A). These quasi-periodic RT jumps should lead to perturbations at 2.5, 5, and 12.5 Hz in the detrended RT time series respectively for the 20-, 10-, and 4-SOA conditions, selectively enhancing or reducing specific frequency components. Some of the spectral perturbations predicted by OTEM fits were indeed observed in the data, most evident in the following two phenomena (Figure 8). First, there was a second or extended peak for the incongruent around 3–5 Hz in the 20- and 10-SOA conditions, respectively echoing the 2.5 and 5 Hz RT jumps. Second and counterintuitively, for the 20-SOA condition, OTEM predicted that the congruent-incongruent phase difference estimated from the detrended RTs should be close to 0 rather than to π, despite attentional oscillations were assumed to be anti-phased. This phase change was due to the vicinity of the frequency of perturbation (2.5 Hz) to the frequency of attentional oscillation (3.3 Hz). Consistent with the OTEM fits, the observed congruent-incongruent phase difference in the 20-SOA condition was close to 0, with the coherence across participants reaching marginal significance. But we are also aware that in the 20- or 10-SOA condition, spectral analysis would hardly allow us to separate a 3–5 Hz oscillation from the 2.5 or 5 Hz RT “jumps” at the block borders due to their vicinity in frequency, thus the 3–5 Hz peaks observed in these two conditions may not necessarily be evidence for 3–5 Hz oscillations in RT.

In addition, OTEM predicts that sequential effects would arise from the trial-by-trial updating of temporal expectation. When the preceding trials have longer SOAs, the temporal expectation for the current SOA would be longer, thus the preparing speed would be lower, resulting in a longer RT. We tested this prediction by regressing the current RT over the SOAs of the last three trials, with the RTs of the last three trials and the current SOA serving as nuisance covariates. As predicted, the coefficients were significantly greater than zero for the last (*t*-tests, *p*s < 0.001 for all conditions), second last (*t*-tests, *p*s < 0.001 for the 20- and 10-SOA conditions) and even up to the third last SOA (*t*-tests, *ps* < 0.05 for the 40- and 20-SOA conditions), with the patterns of the data closely following those of the model fits (Figure 4).

### OTEM predicts the vanish of priming effects at minimum prime-mask interval

Instead of treating the prime and the mask as one unit for priming (“masked prime”, see Eimer, 1999; Eimer & Schlaghecken, 1998), OTEM assumes that the prime and the mask reset the phases of attentional oscillations separately. As we showed above, this assumption allows OTEM to explain the early positive priming and late negative priming in our temporal context experiment as well as in the masked priming experiment of Huang et al. (2015). In these experiments, however, the same 60-msec interval was used between the prime and the mask, which made them temporally yoked, preventing us from reaching stronger conclusions about their separate influences on priming effects.

To test whether the prime and the mask separately reset attentional oscillations, we performed the Int0-Int60 experiment, where for the same participants the prime-to-mask interval was varied across sessions to be 0 or 60 msec. The other settings were similar to the masked priming experiment of Huang et al. (2015), except that SOAs were sampled from a narrower range (32 different SOAs from 0 to 620 msec in a 20-msec step) so that the priming effects, if any, could be stronger due to reduced temporal uncertainty (as we found in the temporal context experiment). For each of the two conditions (Int0 and Int60), each SOA was repeated for 40 times (20 congruent + 20 incongruent).

If the priming effects in the masked priming task had been caused by the (presumably unconscious) perception of the “masked prime” (e.g. Eimer, 1999; Eimer & Schlaghecken, 1998), the Int0 and Int60 conditions would result in similar priming effects as soon as the masking of the prime is similarly efficient. In contrast, OTEM predicts that priming effects should vary with the prime-to-mask interval. The Int0-Int60 experiment also allowed us to distinguish between the two alternative hypotheses about phase resets (Ding et al., 2016), implemented as the additive OTEM and substitutive OTEM models, whose predictions are similar in the Int60 condition (i.e., positive priming followed by negative priming) but very different in the Int0 condition.

The predictions of the two OTEM models are illustrated in Figure 5A-D. Suppose, as we found earlier, the onsets of the prime and the mask start cycles of attentional oscillations that are anti-phased, respectively favoring the congruent and incongruent. When the prime-to-mask interval is minimum, according to the additive OTEM (Figure 5A), the two oscillations would almost cancel out each other and thus lead to null priming effects. Conversely, the substitutive OTEM predicts that the attentional oscillation induced by the mask would overwrite that of the prime from the very beginning and consequently negative priming effects would dominate all through.

We found that the RT patterns in the Int0 and Int60 conditions were distinctively different for the same participants (*N*=19). In the Int60 condition (Figure 5F), we replicated the findings of the masked priming experiment of Huang et al. (2015): the classic positive priming followed by negative priming, as well as the 3–5 Hz oscillated priming (Figure 8A). We further quantified the priming effects using the same time windows as we used in the temporal context experiment (i.e., SOA = 0–60 msec for positive priming and SOA = 100–160 msec for negative priming). For the Int60 condition (Figure 5GH), significant positive priming effects were observed in the previous positive-priming window (*t*-test, *t*(18) = 4.95, p < 0.001) and significant negative priming effects in the previous negative-priming window (*t*-test, *t*(18) = –5.50, p < 0.0001). In contrast, in the Int0 condition (Figure 5E), little priming effects were observed at any SOA and the priming effects quantified in neither time window reached significance (Figure 5GH, *t*-tests, *t*(18) = 0.47, *p* = 0.65, and *t*(18) = 1.79, p = 0.091). The differences between the Int60 and Int0 conditions in the magnitudes of priming effects in the two windows were significant (*t*-tests, *t*(18) = 3.21, p = 0.005, and *t*(18) = –5.17, p < 0.001).

The vanish of priming effects at minimum prime-mask interval is predicted by the additive OTEM (Figure 5AB) but not by the substitutive OTEM (Figure 5CD) or a “masked prime”. For each participant, we further fit both the additive OTEM and substitutive OTEM models to the RT data using maximum likelihood estimates. The “OTEM” in Figure 5EF refers to the additive OTEM fits, whose patterns agree well with the observed RT time series. Similarly, according to a direct comparison of the observed priming effects (Figure 5GH) with the two OTEM fits (Figure 5I–J), the additive OTEM fits successfully captured the observed patterns in both the Int0 and Int60 conditions, while the substitutive OTEM fits failed and apparently mixed up the two conditions.

Figure 5O summarizes the model comparison results between the additive OTEM and the substitutive OTEM for all the masked priming experiments reported in the present paper. (The two OTEMs are identical for unmasked priming.) The Akaike information criterion corrected for small samples (AICc; Akaike, 1974; Hurvich & Tsai, 1989) was used as the metric of goodness-of-fit and smaller AICc indicates better fits. The ΔAICc of a specific model for a specific participant was defined as the difference between the AICc of the model and the participant’s lowest AICc. The protected exceedance probability (Rigoux et al., 2014; Stephan et al., 2009)—the probability a specific model outperforms all the other models—was calculated as the group-level measure for model comparison. In all the experiments, the additive OTEM outperformed the substitutive OTEM in goodness-of-fit (lower summed ΔAICc across participants), though the differences were small expect in the Int0-Int60 experiment, for reasons we stated earlier. In the Int0-Int60 experiment, the AICc difference between the two OTEM models was large and the probability for the additive OTEM to be the winning model (protected exceedance probability) was 96.6%.

It is noteworthy that a common set of phase parameters were estimated for the Int0 and Int60 conditions when the OTEM models were fitted to data. The estimated phase parameters of the additive OTEM (Figure 5MN) were significantly coherent across participants for both the prime (Rayleigh test, *r* = 0.73, *z* = 10.15, *p* < 0.0001) and the mask (Rayleigh test, *r* = 0.61, *z* = 7.04, *p* <0.001). Consistent with our findings in previous experiments and with the phase parameters we had used to generate the predictions in Figure 5AB, the average phases induced by the prime and the mask were respectively close to 0 and π (in sine).

### OTEM outperforms alternative models

We considered two alternative computational models that can produce visual priming effects. Among them, the “constant accumulation model” was based on the accumulator model of Vorberg et al. (2003). The original model, applying only to unmasked priming tasks to explain positive priming, assumes that the prime triggers a constant-speed accumulation of a decision variable towards the congruent. In the constant accumulation model, we extended the model to masked priming tasks with the additional assumption that the mask would change the direction of the constant accumulation towards the incongruent, so that it can produce negative as well as positive priming effects.

A second alternative model was motivated by a widely-accepted idea in the foreperiod task (Luce, 1986; Niemi & Näätänen, 1981; Nobre et al., 2007) that RT for a delayed target depends on the urgency of response at target onset: The higher the urgency, the shorter the RT. In particular, urgency is defined by hazard rate, that is, the conditional probability the target is expected to occur at the next moment, given that it has not occurred yet. To account for oscillated priming, attentional oscillation is assumed as in OTEM. We call the model “oscillated urgency model”, whose assumptions are otherwise the same as OTEM except for how temporal expectation influences RT: The preparedness for the congruent or incongruent is proportional to the instantaneous hazard rate instead of being accumulated over time (see Materials and Methods).

We fit the constant accumulation model and oscillated urgency model to the RT data of Huang et al. (2015) and our two new experiments using maximum likelihood estimates, in the same procedure as we fit OTEM. For all the datasets (Figure 6B), OTEM outperformed the other two models in goodness-of-fit (measured by AICc, smaller is better). According to the group-level Bayesian model selection (Rigoux et al., 2014; Stephan et al., 2009), the probability for OTEM to be the best model (protected exceedance probability) approached 100%.

The deviation of the constant accumulation model from data was obvious (Figure 6A, right column): By definition, it could only predict two opposing RT trends for the congruent and the incongruent—if one decreases with SOA, the other must increase with SOA, a unique finding of Vorberg et al.’s (2003). In the case both the congruent and incongruent RTs decrease with SOA, as in our datasets and most previous visual priming studies (Eimer & Schlaghecken, 2003; Lleras & Enns, 2004; Sumner, 2007), the constant accumulation model would simply fail. The problem with the oscillated urgency model was its inability to produce an enduring positive or negative priming effect (Figure 6A, central column), though its assumptions of attentional oscillation and temporal expectation are similar to those of OTEM. It is because the emergence of positive and negative priming in OTEM relies on the accumulation of preparedness over time as well as on attentional oscillation and temporal expectation. Without the accumulation, the congruent and incongruent RTs would merely follow the fluctuation of attentional oscillations.

### Extended OTEM explains error rates and atypical priming effects

The OTEM introduced above only applies to correct responses, which make up approximately 95% of the trials of most visual priming experiments including ours (Eimer, 1999; Eimer & Schlaghecken, 1998; Huang et al., 2015; Schlaghecken & Eimer, 2002; Verleger, Jaśkowski, Aydemir, van der Lubbe, & Groen, 2004). The error rates, though small, could be patterned (Figure 7A): In the 4-SOA condition, for example, the error rate of the incongruent was higher than that of the congruent around the peak of positive priming (0–60 msec), while it was the reverse around the peak of negative priming (100–160 msec). This cannot be explained by speed-accuracy trade-off, since the accuracy for fast responses was even higher than that of slow responses. Such patterns—positive priming is accompanied by a higher error rate for the incongruent and negative priming by a higher error rate for the congruent—were also observed in previous studies using visual priming tasks (Eimer, 1999; Eimer & Schlaghecken, 1998; Vorberg et al., 2003).

To model error responses, we extended OTEM to include an additional decision process after target onset, where the congruent and incongruent accumulators race against and inhibit each other (see Materials and Methods). Consistent with the observed patterns, the extended OTEM predicts a higher error rate for the incongruent than the congruent at positive priming and the reverse pattern at negative priming (Figure 7B). The RTs and error rates are not cause and effect, but the effects of a common cause: When one accumulator has achieved a higher preparedness than the other at target onset, the subsequent decision process would be biased towards the leading accumulator. In an unmasked priming task, Vorberg et al. (2003) found that the congruent RT decreased with SOA but the incongruent RT increased with SOA, which deviated from the typical finding that both the congruent and incongruent RTs decrease with SOA (e.g. Figure 1B). They also found that error rate increased with SOA. We noticed, however, the SOAs they used were unusually short (no longer than 108 msec), so did all the following studies that had found similar RT patterns (Mattler & Palmer, 2012; Sack, van der Mark, Schuhmann, Schwarzbach, & Goebel, 2009). Such short SOAs would be within the first half cycle of the attentional oscillation of OTEM (approximately 300 msec per cycle), where the attentional gain for the incongruent is still low. That is, the increase of RT with SOA for the incongruent and the decrease of RT with SOA for the congruent they observed can be part of oscillated priming. Using the same SOAs as in Vorberg et al. (2003), the extended OTEM can reproduce their RT and error rate patterns (Figure 7C).

### Additional evidence for the 3–5 Hz oscillated priming

Given that the oscillated priming effect was a major motive of our work, we need to confirm that the 3–5 Hz oscillation reported in Huang et al. (2015) was not just an artifact from spectral analysis. In fact, Huang et al. (2015) reported an extra visual priming experiment (referred as “single-subject experiment”) where each subject completed several thousands of trials so that the subject’s mean RT at each SOA could be reliably estimated (64 trials for each data point). Six subjects participated in this extensive test. As shown in Figure 4 of Huang et al. (2015), “sawtooth” priming patterns (the difference between incongruent and congruent goes up, then down, then up, then down, as a function of SOA) were observed in the raw RT data of all the individual subjects as well as on the group level. The “single-subject experiment” thus provides independent evidence for the 3–5 Hz oscillated priming without using any spectral analysis.

It was noted by a previous reviewer that the specific moving-window detrending procedure used by Huang et al. (2015), with its edge artifact, might cause illusory 3–5 Hz oscillations in the detrended curve when it was applied to a simulated exponentially-decreasing RT curve without any oscillations. To avoid this edge artifact, we used exponential curve for detrending instead in the present study. We have replicated the 3–5 Hz oscillated priming in our two new experiments as well as in the unmasked and masked priming experiments of Huang et al. (2015).

Figure 8 summarizes the results of spectral analysis (amplitude spectrum and the congruent-incongruent phase difference in the 3–5 Hz oscillation) for all the experiments, for which, again, OTEM fits agree well with the data. As we explained in detail earlier for the temporal context experiment, the phase difference in the spectral analysis of the 20-SOA, 10-SOA, or 4-SOA conditions deviated from π partly due to the disturbance of the SOA blocks (i.e. RT “jumps” at block borders). As predicted by OTEM, the phase difference in the Int0 condition was close to 0 and in the other four conditions (unmasked, masked, 40-SOA and Int60) was close to π. The coherence of the phase difference was significant for the Int60 condition (Rayleigh test, *r* = 0.43, *z* = 3.51, *p* = 0.028) but did not reach significance in the unmasked, masked, or 40-SOA condition. We conjecture that the latter three conditions had a lower statistical power than the Int60 condition: each combination of SOA and congruency was repeated for 20 times in the Int60 condition, but for only 16, 16, and 12 times in the unmasked, masked, and 40-SOA conditions. To improve statistical power, we performed spectral analysis for participants pooled from the masked condition and its two replications—the 40-SOA and the Int60 conditions (*N*=53 in total). The phase difference of the aggregated dataset not only was close to π on average but also had highly significant coherence (Rayleigh test, *r* = 0.35, *z* = 6.26, *p* = 0.002).

That OTEM well fit all these datasets itself provides an additional line of evidence for oscillated priming, given that anti-phased attentional oscillations are assumed for the congruent and incongruent. Moreover, the phase reset parameters estimated from different experiments were highly consistent, with the prime and the mask respectively resetting the phases of attentional oscillations to approximately 0 and π (Figures 1I-K, 3D, and 5MN). That is, for the temporal context experiment, similar attentional oscillations were underlying different temporal contexts. As shown in the Int0-Int60 experiment, the same attentional oscillations can also predict the vanish of priming effects at minimum prime-mask interval.

For the model fitting results reported above, we had fixed the frequency of attentional oscillation to be 3.3 Hz, the observed frequency of oscillated priming in spectral analysis. The RTs predicted by OTEM exhibits a similar frequency of oscillated priming as that of the assumed attentional oscillation (Figure 8).

We also tested whether the frequency of attentional oscillation is 3.3 Hz and whether attentional oscillations are necessary at all. To do this, we varied the frequency of attentional oscillation as a free parameter, fit the OTEM model for each frequency, and compared their goodness-of-fit in AICc. Because of computational difficulties, we only considered a limited set of frequencies spanning 0 to 16 Hz. An oscillatory frequency of 0 Hz means no attentional oscillation at all, that is, the prime or mask induces exponentially decreasing or increasing attentional gains. For all the experiments, the best fit occurred at 3.3 or 4 Hz, which outperformed lower or higher frequencies including 0 Hz (Figure 9). On one hand, this justifies our presumption of the 3.3 Hz attentional oscillation as well as our introduction of attentional oscillation as one of the OTEM assumptions. On the other hand, it provides further evidence for the 3–5 Hz oscillated priming.

**Figure 9.**
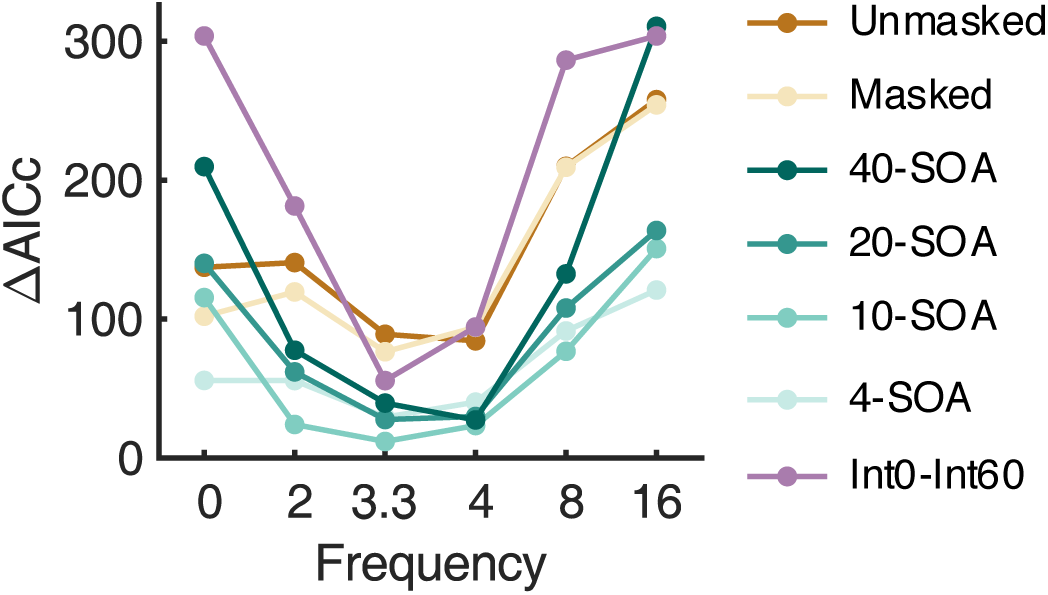
Goodness-of-fit of OTEM as a function of the assumed attentional oscillation frequency. For the model fitting results presented in the text and the other plots, the frequency of attentional oscillation was fixed to 3.3 Hz. Here we compared the model fitting results that were based on different frequencies including 3.3 Hz. Smaller ΔAICc indicates better goodness-of-fit. Curves in different colors denote different experimental conditions (notations same as Figure 8). In all the experiments and conditions, the best fit occurred at 3.3 or 4 Hz, which outperformed lower or higher frequencies including no oscillations (0 Hz).

### Summary of results

To summarize, we developed OTEM as a computational model that combines attentional oscillation and temporal expectation to explain visual priming effects in RT and tested it in four human behavioral experiments. OTEM accurately reproduces all three kinds of priming effects—positive priming, negative priming, and oscillated priming, while the alternative models fail to do so. It correctly predicts temporal expectation and attentional oscillation would influence visual priming effects. It explains not only the averaged priming effects, but also trial-by-trial variations in RT. Most surprisingly, OTEM shows that the classic, sustained positive and negative priming effects can be emergent properties of attentional oscillation and temporal expectation, a prediction that qualitatively holds for a wide range of parameter values.

By explaining the classic priming effects as emergent properties of OTEM, we are not at any position doubting the ecological significance of positive or negative priming. Emergent properties of a system can be important by themselves, just like consciousness is believed to be an emergent property of the brain (Minsky, 1986).

Our findings raise a general concern that the results of priming experiments should be interpreted in their temporal contexts other than by individual SOAs. Some of the previously found differences between priming experiments at the same SOAs might be explained away by temporal contexts. A similar concern is raised for the interval between the prime and the mask, which is not necessarily related to whether the perception of the prime is conscious.

## Discussion

### Relation to previous theories of visual priming effects

What causes visual priming effects? Theories with incremental assumptions about a perceptual or motor bias have been proposed to explain the increasingly richer phenomena. Positive priming had been assumed to be triggered by the prime to favor the congruent (Damian, 2001; Eimer, 1999; Eimer & Schlaghecken, 1998; Neumann & Klotz, 1994; Vorberg et al., 2003). When negative priming was later found in masked priming tasks, self-inhibition of the prime (Bowman et al., 2006; Eimer & Schlaghecken, 2002; Schlaghecken & Eimer, 2006; Schlaghecken, Rowley, Sembi, Simmons, & Whitcomb, 2007) or bias reversed by the mask (Lleras & Enns, 2004; Verleger et al., 2004) was introduced to explain the transition from positive to negative priming with increasing SOA. Efforts were also made to reproduce positive and negative priming in computational models (Vorberg et al., 2003), neural network models (Bowman et al., 2006; Sohrabi & West, 2009), or computational models with biologically plausible neural implementations (Huber & O’Reilly, 2003).

The model we propose here does not necessarily conflict with previous theories. Neither is OTEM intended to be an exclusive explanation for priming effects that rules out alternative accounts such as perceptual aftereffect (Fritsche, Mostert, & de Lange, 2017; Klauer & Dittrich, 2010).

Almost all previous theories considered the visual priming effects as passively determined by the stimuli. Instead, OTEM implements the idea of active sensing (Lakatos et al., 2008, 2013, 2009; Schroeder, Wilson, Radman, Scharfman, & Lakatos, 2010; Zion Golumbic et al., 2013) and assumes that the perceptual system (1) rhythmically samples the congruent and incongruent, and (2) adaptively prepares for the prospective targets. This novel combination of attentional oscillation with temporal expectation enables OTEM to accommodate the recent empirical finding of oscillated priming as well as the classic effects.

Prior to OTEM, temporal expectation was hardly considered in the literature of priming effects, though it has received growing attention in perception and memory (Nobre & van Ede, 2017). Consequently, previous priming theories might explain the decrease of RTs with increasing SOA (e.g. the nROUSE model of Huber and O’Reilly, 2003) but not the RT “jumps” with increasing SOA (Figure 3A). In contrast, with the introduction of temporal expectation that distinguishes SOA and temporal uncertainty, OTEM predicts both the observations.

OTEM also predicts that the classic priming effects may change with temporal context, since temporal expectations are learned from recent experience. As found in the temporal context experiment, the magnitudes of positive and negative priming effects differed across the four temporal expectation conditions: higher temporal uncertainty led to smaller priming effects (Figure 3BC). Similarly, Naccache, Blandin, and Dehaene (2002) found that the classic priming effects would vanish when the arrival time of the target was barely predictable.

With all three priming effects arising from a common process, OTEM also naturally predicts that positive and negative priming are closely related. In agreement with this prediction, Boy and Sumner (2010) showed that both positive and negative priming effects are proportional to the strength of stimulus-response associations: When a novel stimulus-response mapping is used, it takes time to build associations, during which positive and negative priming grow with time at the same pace.

### Neurobiological motivations for OTEM

OTEM is intended to capture the computational heuristics of the neural computations underlying visual priming effects. The neurobiological motivations for some of its key assumptions, such as attentional oscillation (VanRullen, 2013, 2016) and temporal expectation (de Lange et al., 2013; McGuire & Kable, 2013), have been specified in the Introduction. We describe the others below.

There have been elegant models (Huber & O’Reilly, 2003) on how positive priming may spontaneously transition into negative priming with increased exposure to the prime. For the visual (arrow) priming tasks we treated here, however, negative priming can hardly be explained solely as an effect of the prime, because negative priming effect has seldom been observed in unmasked priming tasks (Huang et al., 2015; Jaśkowski et al., 2008; Klotz and Wolff, 1995; Vorberg et al., 2003; but see Klauer and Dittrich, 2010 for an exception). Moreover, it has been found that negative priming would be weaker (Schlaghecken & Eimer, 2006) or even be absent (Jaśkowski & Przekoracka-Krawczyk, 2005; Lleras & Enns, 2006; Verleger et al., 2004) unless the mask shares visual features with the prime (see the arrow-shaped mask in Figure 1E for an example). In other words, masks are not passive stimuli that simply render the prime invisible but play an active role in the priming effects.

A mask that contains features of the prime, according to Huber (2008), will induce neural habituation to the prime and highlight the opposite features. Similar ideas have been expressed by Lleras and Enns (2004) in the object updating theory and are analogous to perceptual aftereffect (Webster, 2015). We therefore assume that the mask, like the prime, resets the attentional oscillation but to a different phase that favors more of the opposite features. To capture potential individual differences, the exact phase reset by the mask is left to be a free parameter in OTEM, which turned out to be highly consistent in all our masked priming experiments and was anti-phased to that of the prime. That the prime and the mask induce anti-phased attentional oscillations predicts the vanish of priming effects for minimum prime-mask interval.

Temporal discounting has been found in human perception and action (Shadmehr, Orban de Xivry, Xu-Wilson, & Shih, 2010) as well as in economic decision-making (Frederick et al., 2002; Kable & Glimcher, 2007). In OTEM, we introduced temporal discounting to characterize the tendency to prepare more intensely for a near-future target compared to a far-future target that is equally likely to occur. When the discounting rate is zero, any future event would be treated as urgent as the present event and temporal expectation would thus be meaningless. On the other extreme, when the discounting rate is infinity, only the present matters, which is effectively the assumption adopted by many previous models including but not limited to the hazard rate model of temporal expectation (Los et al., 2014; Los & Van Den Heuvel, 2001; Luce, 1986; Niemi & Näätänen, 1981; Nobre et al., 2007).

In the presence of temporal uncertainty, the probability of a future event can leak into the present such that the participant would appear to prepare for the future even if she only cares about the hazard rate at the current moment. At first sight, temporal uncertainty seems to play a role interchangeable with temporal discounting, which renders the latter redundant. However, a model with the former alone would predict an increase of RT with increasing temporal uncertainty, other things being the same (Tsunoda & Kakei, 2011), which is opposite to our empirical finding that, in the 20-, 10-, or 4-SOA condition, blocks with longer SOAs tend to have shorter RTs (Figure 3A), though longer SOAs correspond to larger temporal uncertainty due to Weber’s law (Allan, 2001; Gibbon, 1977). Therefore, for both theoretical and practical reasons, we chose to include both temporal uncertainty and temporal discounting in OTEM.

OTEM assumes that two threads of perceptual or motor preparations develop in parallel before target onset. Simultaneous preparation for multiple potential actions has been reported in neuronal activities (Cisek, 2005, 2007; Cisek & Kalaska, 2010). Such pre-target preparing activities are largely driven by temporal expectation and can bias the subsequent actions (de Lange et al., 2013; Donner, Siegel, Fries, & Engel, 2009). According to OTEM, visual priming effects in RT arise from pre-target preparation instead of post-target decision (but also see the extended OTEM). It echoes the findings that attentional status (Nunez, Vandekerckhove, & Srinivasan, 2017) or temporal expectation (Jepma, Wagenmakers, & Nieuwenhuis, 2012) mostly influences the non-decision time instead of decision time in RT.

### Limitations and future directions

We do not intend to explain all important aspects of visual priming effects, such as how the strength of priming effects is influenced by the features of the prime and target (Huber & O’Reilly, 2003; Kruijne & Meeter, 2017). Neither do we claim that our model provides an implementation-level explanation for visual priming effects. Our work, in Marr’s (1982) term, is between the computational and algorithm levels based on neurobiologically motivated assumptions.

There is evidence that attention can switch rhythmically among more than two targets (VanRullen, Carlson, & Cavanagh, 2007).Though motivated by laboratory tasks with only two potential primes and targets, OTEM can be readily applied to situations where the primes or targets have more than two potential states, by assuming that a prime will trigger attentional oscillations among multiple potential targets. Similarly, OTEM predicts that temporal expectations would modulate the priming effects for multiple primes or targets, a prediction that can be tested in the future.

A coexistence of sustained and oscillated effects has also been found in spatial cueing tasks. More generally, OTEM offers a new perspective that may unify the attention-related classic findings with the increasing evidence for attentional oscillations in visual perception. One future question is, may other attention-related classic phenomena be explained in a similar framework?

## Acknowledgments

MW was supported by Key Laboratory of Machine Perception Collaborative Research Grant K-2019-04. YH was supported by Shenzhen Science and Technology Research Funding Program JCYJ20170818161400180 and Guangdong Provincial Key Laboratory of Brain Connectome and Behavior 2017B030301017. HL was supported by National Natural Science Foundation of China grant 31930052. HZ was supported by National Natural Science Foundation of China grants 31871101 and 31571117, and funding from Peking-Tsinghua Center for Life Sciences. Part of the analysis was performed on the High-Performance Computing Platform of the Center for Life Sciences at Peking University and supported by the High-performance Computing Platform of Peking University. The funders had no role in study design, data collection and analysis, decision to publish, or preparation of the manuscript.

## Notes

#### Summary of Updates

The most important revision is to include the results of a new visual priming experiment, where we tested an additional key prediction of our computational model: The interval between the prime and the mask would influence priming effects. The results are consistent with the predictions of our model.

